# Longitudinal profiling of the macaque vaginal microbiome reveals similarities to diverse human vaginal communities: implications for use as a pre-clinical model for bacterial vaginosis

**DOI:** 10.1101/2020.12.14.422805

**Authors:** Nicholas S. Rhoades, Sara M. Hendrickson, Danielle R. Gerken, Kassandra Martinez, Ov D. Slayden, Mark K. Slifka, Ilhem Messaoudi

**Affiliations:** Department of Molecular biology and Biochemistry, University of California Irvine, Irvine, CA, USA; Division of Neuroscience, Oregon National Primate Research Center, Beaverton OR, USA; Division of Reproductive & Developmental Sciences, Oregon National Primate Research Center, Beaverton, OR, USA

## Abstract

The vaginal microbiota plays an important role in women’s reproductive and urogenital health. Disturbances in this microbial community can lead to several adverse outcomes including pelvic inflammatory disease, bacterial vaginosis (BV) as well as increased susceptibility to sexually transmitted infections, miscarriage, and pre-term births. It is now well accepted that while the microbiome of healthy women in the developed world is dominated by *Lactobacillus* species, vaginal communities in asymptomatic women, especially those in the developing world, can be comprised of a diverse set of micro-organisms. The presence of a diverse vaginal microbiome has been associated with increased susceptibility to HIV infection but their implications for women’s health remain poorly understood. Rhesus macaques are an excellent translational animal model due to significant physiological and genetic homology with humans. In this study, we performed a longitudinal analysis of clinical and microbiome data from 16 reproductive age female rhesus macaques. Many animals showed hallmarks of BV, including Nugent scores above 7 and high vaginal pH. At both the taxonomic and functional level, the rhesus macaque vaginal microbiome was most similar to that of women who harbor a diverse vaginal community associated with asymptomatic/symptomatic bacterial vaginosis. Specifically, rhesus macaque vaginal microbiomes harbored a diverse set of anaerobic gram-negative bacteria, including; *Snethia*, *Prevotella, Porphyromonas*, and *Mobilluncus*. Interestingly, some animals were transiently colonized by *Lactobacillus* and some with *Gardnerella*. Our in-depth and comprehensive analysis highlights the importance of the model to test interventions for manipulating the vaginal microbiome.

**IMPORTANCE:** It is widely accepted that the “healthy” vaginal microbiome of the majority of women in the developed world is dominated by *Lactobacillus* species. However, in the developing world, a majority of women are colonized by diverse microbial communities, typically associated with bacterial vaginosis, but remain asymptomatic. Many questions remain about the drivers of this disparity and potential interventions to alter the vaginal microbiome. Rhesus macaques provide an excellent translational model due to significant physiological and genetic homology with humans. In this study, we performed a longitudinal analysis of clinical and microbiome data from a large cohort of reproductive age rhesus macaques. At the taxonomic, genomic, and functional level, the rhesus macaque vaginal microbiome was most similar to that of humans who harbor a diverse vaginal community associated with asymptomatic/symptomatic bacterial vaginosis. Our in-depth and comprehensive analysis highlights the utility of macaques to test interventions for manipulating the vaginal microbiome.

## INTRODUCTION

The vaginal microbial community modulates several critical physiological functions and protect the host from infection with pathogenic organisms. The “healthy” vaginal microbiome (VM) in women in the developed world is typically dominated by *Lactobacillus* species, with *L. crispatis*, *L. iners*, *L. johnsonii*, and *L. gasseri*. These microbes are considered keystone microbes that rarely co-occur in individuals (*1, 2*). *Lactobacilli* inhibit the growth of other vaginal microbes via the production of lactic acid through fermentation which decreases the pH of the vaginal environment (pH 3.5-5.5) inhibiting the growth of other microbes (*3*). *Lactobacillus* species also competitively exclude genitourinary pathogens thereby preventing infection. Disruptions of this community have significant implications for women’s health (*4–6*). However, approximately ~20% of women in the developed world and ~50% in the developing world harbor a diverse VM with reduced abundance or absence of *Lactobacilli* and a high abundance of taxa such as *Gardnerella*, *Prevotella*, *Snethia*, and *Mobiluncus* (*1, 4*). This diverse community is often associated with a heightened inflammatory state, increased susceptibility to sexually transmitted diseases notably HIV, and the development of bacterial vaginosis (BV) (*7*).

BV is an inflammatory disorder and the most common vaginal infection in the US affecting ~ 30% of women ages 15-44 (*8*). However, the prevalence of BV can vary widely based on geographic location and racial background (*8*). BV is caused by bacterial overgrowth, disruption of the commensal microbial community, and a pro-inflammatory environment (*9*). While typically asymptomatic, BV can result in discomfort and more importantly increased susceptibility to sexually transmitted diseases, preterm birth, and pelvic inflammatory disease (*6, 10, 11*). The antibiotics Metronidazole and/or Clindamycin are the current standard treatment for BV. However, BV recurs in ~50% of women within 12 months of antibiotic treatment (*12*). Additionally, the production of biofilms and the development of other antimicrobial resistance mechanisms will most likely reduce the effectiveness of antibiotics treatments over time (*13, 14*). Recently, VM transplants have been shown to be effective in the treatment of intractable BV (*15*). However, microbiome transfer procedures have recently come under scrutiny due to safety concerns (*16, 17*).

Another potential intervention is the use of pre-biotics to shift a microbial community to a more “beneficial” state. Pre-biotics are molecules and/or nutrients that are metabolically accessible to microbes with the goal of enriching for beneficial microbes and/or depleting undesired microbes. An example of such an intervention is the consumption or supplementation of dietary fiber to enrich the gut microbiome in beneficial short-chain fatty acid-producing microbes (*18, 19*). The use of pre-biotics has also been proposed as a cheap and easily accessible alternative to antibiotic treatment for BV (*20, 21*). Of interest is the intravaginal application of di- or polysaccharides that are preferentially metabolized and fermented by *Lactobacillus* to produce lactic acid and hydrogen peroxide, thereby generating a low pH vaginal environment and inhibiting the growth of BV-associated bacteria. Clinical trials are currently underway to test the use of intravaginal Lactose (NTC03878511), Glucose (NCT03357666), and Glycogen (NCT02042287) to treat BV symptoms. Sucrose in particular has shown potential for improving clinical markers of BV in humans (*22*) and shifting the VM of rhesus macaques (*23*). However, the clinical study only examined changes in clinical symptoms and Amsel criteria at one-time point after 14 days of treatment and did not interrogate changes in microbial communities (*22*). The preclinical study in macaques utilized animals with a vaginal microbial community that had >1% *Lactobacillus;* did not identify the *Lactobacillus* species present; nor did it report individual animal data (*23*). Therefore, it is still unclear whether a sucrose intervention could improve clinical outcomes when the frequency of *Lactobacillus* species is extremely low, which *Lactobacillus* strains can respond to sucrose treatment, and how effective is this intervention.

To address these questions, we utilize a combination of clinical tests, 16S rRNA amplicon sequencing, and shotgun metagenomics to characterize the taxonomic and functional landscape of the rhesus macaque VM before and after intra-vaginal sucrose treatment. Additionally, we determined the relatedness of rhesus macaque and human vaginal microbes with whole-genome resolution. Previous studies have defined the taxonomic landscape of pigtail (*24*), cynomologus (*25*), and rhesus (*26*) macaques VM. Together these studies have shown that macaques harbor a diverse vaginal community (*24–26*) that shares taxa with the diverse community state type associated with BV in humans (*25*). These patterns include a low abundance of *Lactobacillus spp.* and a high abundance of *Snethia*, *Prevotella*, and *Mobiluncus* among others. However, these previous studies have been limited to amplicon sequencing techniques that are limited in resolution and did not address the functional potential of the macaque vaginal community or examine this similarity at a whole-genome level. The data presented herein further strengthen the case for using macaques to test interventions for altering the VM.

## METHODS

### Sample collection and cohort information

All macaque studies were reviewed and approved by the OHSU/ONPRC Institutional Animal Care and Use Committees (IACUC). The animals were socially housed indoors at the Oregon National Primate Research Center (ONPRC) following standards established by the US Federal Animal Welfare Act and *The National Research Council Guide for the Care and Use of Laboratory Animals*. All animals were tested annually for simian viruses (Simian Immunodeficiency Virus, Simian Retrovirus 2, Macacae herpesvirus 1, and Simian T lymphotropic virus) and received a mammalian old tuberculin test semi-annually. The monkeys underwent pre-assignment evaluations prior to initiation of the study. Qualifications of assignment to the study included normal menstrual cycle, age 5-8 years, and a healthy weight of 4-8kgs, and all had to be void of gastrointestinal issues and antibiotic use for greater than three months. At the pre-assignment screening, Nugent score and microbiome samples were collected.

Samples were collected for 16S rRNA amplicon sequencing, metagenomics, and Nugent scoring. Samples used for Nugent scoring, including pH analysis and whiff test, were collected using polyester swabs (Fisher Scientific; item 23-400-122). Samples used for microbiome analysis were collected with Copan Swabs (Fisher Scientific; 23-600-957) and stored in 20% glycerol (shotgun metagenomics) or immediately snap-frozen upon collection and stored at −80°C until DNA extraction (16S rRNA amplicon sequencing). To collect the vaginal and rectal swab samples, animals were sedated and placed in a ventral recumbency with their pelvis slightly elevated. A sterile nasal speculum was inserted vaginally to ensure a mid-vaginal sample collection without rectal or vaginal introitus contamination. Preliminary screening identified 3/20 (15%) of adult rhesus macaque females with vaginal pH = 4, absence of Clue cells, and negative Whiff test. These animals were not included in the longitudinal study. The animals chosen for further study had high pH (pH >5) and other characteristics of vaginal microbiome dysbiosis.

To minimize the natural variability that may occur over the menstrual cycle, animals received 21 days of combination oral contraceptives (Portia; Teva Pharmaceuticals USA, Inc.) to synchronize the group. One day after the last dose of oral contraceptives, samples were collected (Nugent scores, Clue cells, Whiff test, microbiome, and metagenomics), and animals menstruated approximately 24-48 hours later. Next, animals underwent post-menstruation sample collection (Nugent scores, Clue cells, Whiff test microbiome, metagenomics), and then began five days of gel treatment administration.

Animals were randomly designated to one of two treatment groups, Control (vehicle only; n=8) or Sucrose (10% sucrose; n=8). The sucrose treatment consisted of 1.6% xanthan gum (Xantural 180; CP Kelco) added to a 10% sucrose solution (Sigma Aldrich) to create a gel property and brought to a pH of 5.0 using lactic acid (Fisher Scientific). The control treatment did not contain sucrose. Vaginal administration of the gel occurred daily using a 5ml slip-tip syringe, with control animals received the xanthan gum gel only.

### Hormone quantification

Serum concentrations of estradiol (E_2_) and progesterone (P_4_) were assayed by the ONPRC Endocrine Technologies Core. Hormone concentrations were determined using a chemiluminescence-based automated clinical platform (Roche Diagnostics Cobas e411, Indianapolis, IN). Serum E2 and P4 assay ranges were between 5-3000 pg/ml and 0.05-60 ng/ml, respectively. E_2_ and P_4_ intra-assay CVs were 8.3 and 4.2%, whereas inter-assay CVs were 5.9 and 6.3%, respectively.

### Clinical data generation

Nugent’s scoring was carried out by a trained microbiologist with 4 years of experience performing Nugent scores in a CLIA-approved clinical laboratory using established criteria for clinical studies (*27*). To complement Nugent scoring, three of four Amsel criteria (*28*) were also assessed including vaginal pH, presence of Clue cells, and Whiff test. Vaginal discharge, the fourth Amsel criteria, was not measured. The presence of Clue cells (i.e., vaginal epithelial cells coated with bacteria, resembling a “sandy” appearance) was determined and samples with ≥20% Clue cells were considered positive for this test. Whiff tests were performed by adding 10% KOH to a sample of vaginal fluid and the presence of fishy odor was interpreted as a positive test while its absence was determined as a negative test result. A full breakdown of clinical measurements can be found in supplemental table 1 **(Supp. Table 1)**.

### 16S rRNA gene library construction and sequencing

Total DNA was extracted from vaginal swabs using the Blood and Tissue DNA Isolation KIT (MO BIO Laboratories, Carlsbad, CA, USA). Rectal swabs were extracted using the PowerSoil DNA Isolation KIT (MO BIO Laboratories, Carlsbad, CA, USA). This DNA was used as the template to amplify the hypervariable V4 region of the 16S rRNA gene using PCR primers (515F/926R with the forward primer containing a 12-bp barcode) in duplicate reactions containing: 12.5 ul GoTaq master mix, 9.5 ul nuclease-free H20, 1 ul template DNA, and 1 ul 10uM primer mix. Thermal cycling parameters were 94°C for 3 minutes; 35 cycles of 94°C for 45 seconds, 50°C for 1 minute, and 72°C for 1 minute and 30 seconds; followed by 72°C for 10 minutes. PCR products were purified using a MinElute 96 UF PCR Purification Kit (Qiagen, Valencia, CA, USA). Libraries were sequenced (1 x 300 bases) using Illumina MiSeq.

### 16S rRNA gene sequence processing

Raw FASTQ 16S rRNA gene amplicon sequences were uploaded and processed using the QIIME2 version 2019.10 (*29*) analysis pipeline as we have previously described (*30*). Briefly, sequences were demultiplexed and quality filtered using DADA2 (*31*), which filters chimeric sequences and generated sequence variants were then aligned using the MAFFT (*32*) and a phylogenetic tree was constructed using FastTree2 (*33*). Taxonomy was assigned to sequence variants using q2-feature-classifier against the SILVA database (release 119) (*34*). To prevent sequencing depth bias, samples were rarified to 10,000 sequences per sample before alpha and beta diversity analysis. QIIME 2 was also used to generate the following alpha diversity metrics: richness (as observed taxonomic units), Shannon evenness, and phylogenetic diversity. Beta diversity was estimated in QIIME 2 using weighted and unweighted UniFrac distances (*35*).

Analysis of the unweighted Unifrac distance (based on the presence/absence of microbes) revealed that 9 vaginal samples were very similar to the fecal microbiome (**Supp. Figure 1A**). These 9 samples had a significantly higher number of observed Amplicon sequence Variants (ASVs) than the group average of all other vaginal samples (**Supp. Figure 1B**) and a high relative abundance of taxa typically found in the fecal microbiome (**Supp. Figure 1C**). Therefore, these nine VM samples were removed from future analysis along with eight additional samples that did not meet our minimum sequencing depth threshold of 10,000 reads (**Supp. Figure 1D**).

16S rRNA gene amplicon sequencing data obtained from vaginal swabs from 236 women from Umlazi, South Africa aged 18 to 23 was obtained from Gossman et al. (*4*). These samples were imported into QIIME 2 and rarified to 13,000 reads per sample. Taxonomy was assigned using the full-length SILVA database (release 119) at the 99% OTU cutoff. Genus-level (L6) taxonomy tables were merged, and Bray-Curtis dissimilarity matrices were generated using QIIME 2.

### Shotgun metagenomics

Shotgun metagenomic libraries were prepared for vaginal samples obtained from all animals at the initiation of oral contraceptives using the Illumina Nextera Flex library prep kit, per the manufacturer’s recommended protocol, and sequenced on an Illumina HiSeq 4000 2 × 100. Raw demultiplexed reads were quality filtered using Trimmomatic (*36*), and potential host reads were removed by aligning trimmed reads to the *Macaca mulatta* genome (Mmul 8.0.1) using BowTie2 (*37*). After quality filtering and decontamination, an average of 6.08 million reads (min 0.517, max 30 million reads) per sample were used for downstream analysis. Samples with less than 1 million reads after quality filtering were excluded from downstream analysis. Trimmed and decontaminated reads were then annotated using the HUMAnN2 pipeline using the default setting with the UniRef50 database and assigned to GO terms (*38*). Functional annotations were normalized using copies per million (CPM) reads before statistical analysis (*39–41*).

Trimmed and decontaminated reads were assembled into contigs using meta-SPAdes with default parameters (*42*). Assembled contigs <1kb were also binned into putative metagenomically assembled genomes (MAGs) using MetaBat (*43*). Genome completeness/contamination was tested using CheckM (*44*), and all bins with a completeness > 80% and contamination < 2% were annotated using PATRIC (*45*). The taxonomy of draft genomes was determined using PATRICs’ similar genome finder.

Shotgun metagenomic sequencing data was obtained from Lev-Sagie et.al (*15*) and Oliver et. al (*46*). Human samples were classified as “ Lactobacillus dominated” samples if the relative abundance of *Lactobacillus* >90%. Samples classified as “Diverse human (BV symptomatic)” were the pre-transplant samples from Lev-Sagie et.al (*15*). Samples classified as “Diverse human (asymptomatic)” were obtained from Oliver et. al (*46*) with a *Lactobacillus* relative abundance < 90%. To avoid pseudoreplication only samples from the earliest time-point were used. Sequences were annotated using the HUMAnN2 pipeline as described above for rhesus vaginal samples.

### Statistical analysis

All statistical analyses were conducted used PRISM (V8). QIIME 2 was used to calculate alpha-diversity metrics; observed OTUs, Shannon evenness, and beta diversity; and weighted and unweighted UniFrac distances. Bray-Curtis dissimilarity matrices were constructed for species-level relative abundance. Unpaired t-tests and one-way ANOVAs with post hoc correction were implemented using PRISM (V8) to generate *p* values. The LEfSe algorithm was used to identify differentially abundant taxa and pathways between groups with logarithmic linear discriminant analysis (LDA) score cutoff of 2.

## RESULTS

### Rhesus macaques display the clinical hallmarks of BV and are colonized by a diverse vaginal microbiome

To determine if Rhesus macaques can serve as a model of BV, we screened 9 reproductive age female rhesus macaques. Of these animals, six (66%) had a Nugent score >7 and the presence of Clue cells (Table 1). Additionally, 5 of these 6 animals had a vaginal pH of ≥6.0 (Table 1). These data suggest that the majority of rhesus macaque females display the clinical hallmarks of BV. Profiling of the vaginal microbial (VM) communities using 16S rRNA gene amplicon sequencing showed a high relative abundance of bacterial taxa associated with BV, including; *Snethia*, *Mobiliuncus*, *Prevotella*, and *Gardnerella* (Table 1). Interestingly, 3 animals (33%) included in this screen had a high relative abundance of *Lactobacillus* (43-84%). These data suggest that a subset of RM can naturally harbor a *Lactobacillus* dominated vaginal community.

**Table 1.**

| **Animal** | **1** | **2** | **3** | **4** | **5** | **6** | **7** | **8** | **9** | **10** |
| --- | --- | --- | --- | --- | --- | --- | --- | --- | --- | --- |
| Nugent score | 9 | 9 | 8 | 5 | 10 | 8 | 7 | 2 | 5 | N.C. |
| Clue cells >20% | Yes | Yes | Yes | No | Yes | Yes | Yes | No | No | N.C. |
| Vaginal pH | 7 | 7.5 | 5.5 | 7.5 | 7 | 7.5 | 5.5 | 6 | 4.5 | 8 |
| 16S amplicon relative abundance |  |  |  |  |  |  |  |  |  |  |
| Leptotrichiaceae (Snethia) | 27.26 | 21.26 | 30.52 | 89.78 | 9.7 | 9.37 | 0.36 | 2.78 | 2.13 | 0.05 |
| Lactobacillus | 0.02 | 0 | 0 | 0.13 | 0.03 | 0.03 | 42.98 | 55.75 | 84.44 | 0.1 |
| Prevotella | 14.84 | 5.02 | 13.58 | 0 | 7.46 | 8.41 | 4.97 | 0.25 | 0.02 | 0.02 |
| Mobiluncus | 10.53 | 14.97 | 3.07 | 0 | 7.01 | 8.16 | 0.46 | 0.62 | 0.01 | 0 |
| Gardnerella | 0 | 0 | 0 | 0 | 1.17 | 0 | 0 | 6.01 | 8.29 | 0 |
| Porphyromonas | 6.66 | 10.78 | 11.08 | 0.29 | 2.95 | 5.18 | 0.32 | 0.36 | 0 | 8.7 |
| Trichococcus | 0.31 | 2.18 | 1.1 | 0.18 | 1.18 | 0.62 | 10.76 | 0.67 | 0.01 | 61.01 |
| Campylobacter | 3.74 | 3.79 | 7.1 | 0 | 5.74 | 15.94 | 0.07 | 0.27 | 0 | 0.02 |
| Anaerococcus | 0.43 | 0.07 | 1.75 | 0.35 | 0.27 | 0.37 | 15.77 | 0.75 | 0.11 | 15.1 |
| Catonella | 0 | 0 | 0 | 0 | 8.91 | 9.87 | 0 | 0.14 | 0 | 0 |
| Fusobacterium | 1.71 | 0.3 | 16.22 | 0.02 | 0 | 0 | 0.01 | 0.11 | 0.01 | 0 |
| Peptoniphilus | 4.14 | 3.81 | 1.62 | 0.24 | 1.56 | 3.91 | 0.27 | 0.58 | 0.01 | 1.7 |

N.C. = Not Collected

### Intra-Vaginal Sucrose treatment does not alter the vaginal microbiome of rhesus macaques

We screened an additional 20 animals for inclusion in a longitudinal study to determine the efficacy of vaginal sucrose gel to improve BV clinical indicators and increase the abundance of *Lactobacillus* in the VM. Initial screening of these 20 animals indicated that 4 animals did not display the clinical indicators of BV (pH <5 and/or Nugent score <4) and were excluded from the longitudinal study. The remaining 16 animals were split evenly into two groups. Animals in group 1 received seven daily applications of a sucrose gel vaginally, while animals in group 2 received seven daily applications of the gel alone. Since hormonal levels can influence VM composition (*25, 47, 48*), animals were administered an oral contraceptive for 21 days (1-month prior to sucrose treatment) to synchronize the menstrual cycles of the animals (**Figure 1A**). Contraceptive treatment resulted in a significant decrease in the levels of progesterone and estradiol (**Figure 1 B, C**). Seven days after cessation of the contraceptive treatment, the animals were then treated with either sucrose or placebo gel intravaginally for 7 days. These animals were sampled at 5 time-points across 35 days, to collect clinical (Vaginal pH, Nugent scores, whiff test, and Clue cells), circulating hormone levels (Progesterone and Oestrodiol), and microbiome (16S amplicon sequencing) data (**Figure 1A**).

**Figure 1:**
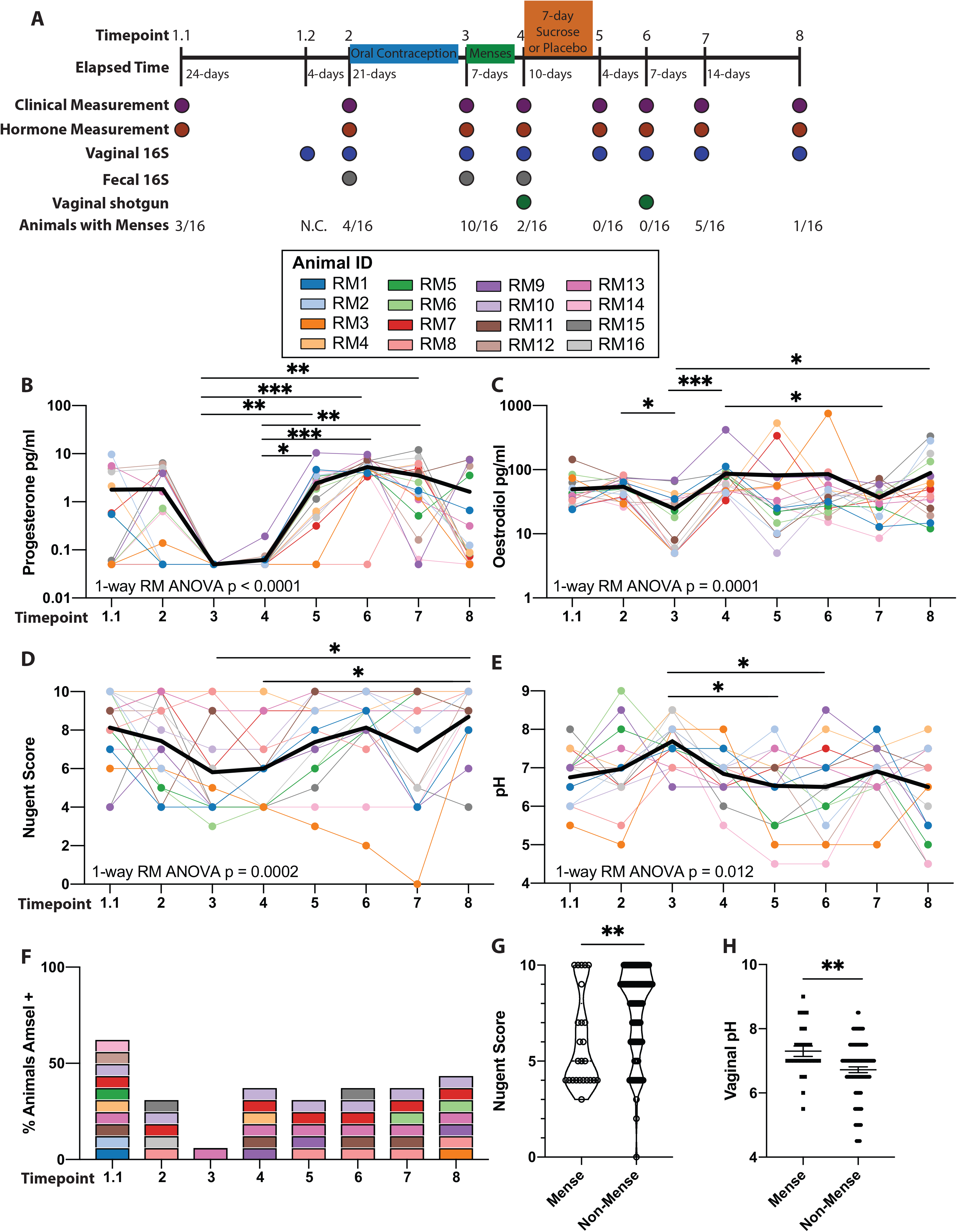
Longitudinal assessment of clinical markers associated with BV. (A) Study design and timeline. (B-E) Scatter plot of measured systemic (B) Progesterone, (C) Oestrodiol, (D) Nugent scores, and (E) pH. Each dot represents an individual sample with solid lines connecting samples from the same individual across time. The bold black line represents the mean value across time. Significance was measured by non-parametric 1-way repeated measures ANOVA (Friedman test) with Dunn’s posthoc comparisons between time points. Overall p-value denoted on each graph and bars denoting significance of post hoc tests, * p < 0.05, ** p < 0.01, *** p < 0.001. (F) Stacked bar graph denoting the percentage of animals that meet Amsel criteria. Colors correspond to those used in panels B-E. (G, H) Scatter plot of (G) Nugent scores and (H) vaginal pH comparing samples collected in animals with and without menses. Significance determined by unpaired t-test, ** p < 0.01.

Sucrose treatment did not alter clinical measurements (pH, Nugent score, Whiff test, or presence of Clue cells) or levels of progesterone or estradiol up to 28 days after treatment (**Supp. Figure 2 A-F**). Additionally, we observed no differences in VM diversity, overall community composition (**Supp. Figure 3 A-C).** Sucrose treatment also did not increase the relative abundance of *Lactobacillus* (**Supp. Figure 3D**). Since sucrose treatment did not lead to measurable changes in clinical or microbiome measurements, we combined the two groups to generate a longitudinal dataset to further our understanding of rhesus macaque VM community dynamics.

Analysis of the clinical markers of BV in all 16 animals showed association with menstrual cycle. Specifically, menses were associated with reduced Nugent scores and a smaller proportion of animals that meet all three measured Amsel criteria while pH levels increased (**Figure 1D-F**). The association between menstrual cycle and these clinical markers is more evident when comparing Nugent scores and vaginal pH in animals with or without menses (**Figure 1 G, H**).

### The taxonomic landscape of the rhesus macaque vaginal microbiome

Using 16S rRNA gene amplicon sequencing, we characterized the taxonomic landscape of the rhesus VM across eight time-points (**Figure 2A**). The rhesus VM is a low diversity community with an average of ~50 ASVs per sample across all time-points (**Figure 2B**). Interestingly, the overall community composition remained stable across time-points in 7 animals (**Figure 2C**). These “Stable” communities were dominated by either *Prevotella*, *Prophymonas,* or *Snethia* and were associated with a higher vaginal pH (**Figure 2 A, C**). The VM of the remaining 9 animals was more variable, transitioning between states and communities dominated by less common microbes such as *Gardnerella*, and *Lactobacillus* (**Figure 2C**). Finally, we found that individual (PERMANOVA: p=0.009) rather than time point (PERMANOVA: p=0.42) was the best predictor of community composition.

We next explored the most abundant taxa within rhesus macaques VM. A small number of samples (13/120) were dominated (>50% relative abundance) by a single microbe (**Figure 2A**), including 9 samples dominated by *Snethia*, 2 by *Gardnerella*, and 4 by *Lactobacillus*. The remaining samples contained communities composed of multiple anaerobic bacteria including *Snethia*, *Porphyromonas*, *Prevotella*, *Fastidiosipila*, *Cantonella*, *Mobiluncus*, and *Atopobium* (**Figure 2A**). Additional taxa found in lower abundance across all samples include *Parvimonas*, *Dialister*, *Fusobacterium*, *Treponema*, *Peptoniphilus*, and *Campylobacter* (**Figure 2A**). The relative abundance of the most abundant 25 taxa did not cluster by time of sample collection or menstruation status; however, samples from some animals with a “Stable” VM did cluster together (**Figure 2E**). Interestingly *Lactobacillus* was detectable in 51 samples and found in a relative abundance above 30% in 5 samples. These five samples came from two monkeys across 4 time-points.

**Figure 2:**
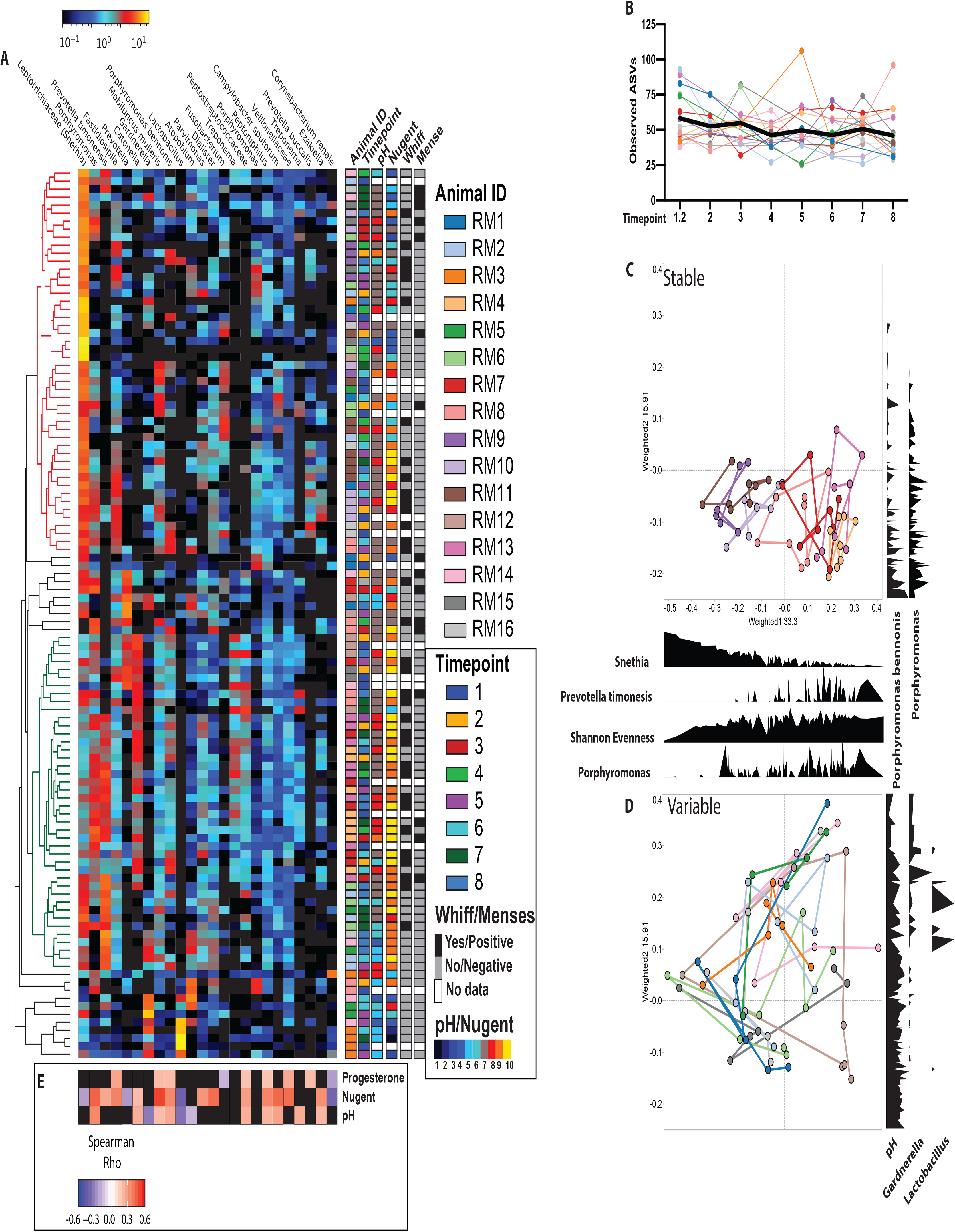
Longitudinal changes in the rhesus macaque vaginal microbiome. (A) Heatmap of the 25 most abundant taxa across all samples ordered from left to right by average abundance. Samples clustered by average linkage of Bray-Curtis distance between samples. Metadata associated with each sample including animal ID, time-point, Nugent score, vaginal pH, whiff test positivity, and menstruation status is provided as well as a heatmap of Spearman rank correlation rho values between vaginal pH, Nugent scores and systemic progesterone levels against the abundance of top 25 microbes. (B) Scatter plot of absolute sequence variants (ASVs) at each time point. Each dot represents an individual sample with solid lines connecting samples from the same individual across time. The bold black line represents the mean value across time. (C, D) Principal coordinate analysis of unweighted UniFrac distance colored by individuals with lines connecting samples collected from the same individual over time, with density plots of key microbial taxa along the PCoA1 and PCoA2 axis.

We next explored the relationship between clinical measures, hormone levels, and the relative abundance of vaginal microbes using Spearman rank correlations (**Figure 2F**). As expected, Nugent score and vaginal pH were positively correlated, while the relative abundance of *Lactobacillus* was negatively correlated with both vaginal pH and Nugent score (**Figure 2F**). Nugent score was positively correlated with systemic Progesterone levels. The relative abundance of *Porphyomonas*, *Mobiluncus mulieris*, and *Campylobacter sputorum* were also positively correlated with Nugent scores (**Figure 2F**). Levels of estradiol did not correlate with clinical markers or relative abundance of specific microbes. Surprisingly *Gardnerella* was also negatively correlated with vaginal pH (**Figure 2F**). However, the vaginal pH of those animals was in the 5-7 range, and those animals harbored a low abundance of *Lactobacillus*.

### The rhesus macaque vaginal microbiome is comparable to human non-*Lactobacillus* dominated vaginal communities

Next, we compared our longitudinal 16S rRNA gene amplicon data to those reported in humans by Gosmann et. al (*4*). This study was selected for the large number of vaginal samples examined and the presence of both *Lactobacillus* dominated (N =83) and diverse communities (N =117). Rhesus macaque samples were divided into “Diverse” (N=105) and “High *Lactobacillus*” (N=5). Due to differences in differences in the methodology, a comparison could only be made qualitatively at the genus level. A principal coordinates analysis shows that the “Diverse” rhesus VM was more similar to “Diverse” than “*Lactobacillus* dominated” human vaginal communities (**Figure 3A, B**). Six bacterial genera were shared between at least one human and one macaque group with ≥1% relative abundance: *Lactobacillus, Prevotella, Gardnerella*, *Snethia, Atopbium,* and *Mobiluncus* (**Figure 3C**). *Megasphera*, *Veillonella*, and Bacteroidales were only found in the diverse human vaginal microbiome (**Figure 3B**). On the other hand, *Porphytomonas, Fastidiosipila, Parvimonas, Dialister, Campylobacter,* and *Catonella* were only found in the “Diverse” macaque VM (**Figure 2B**).

**Figure 3:**
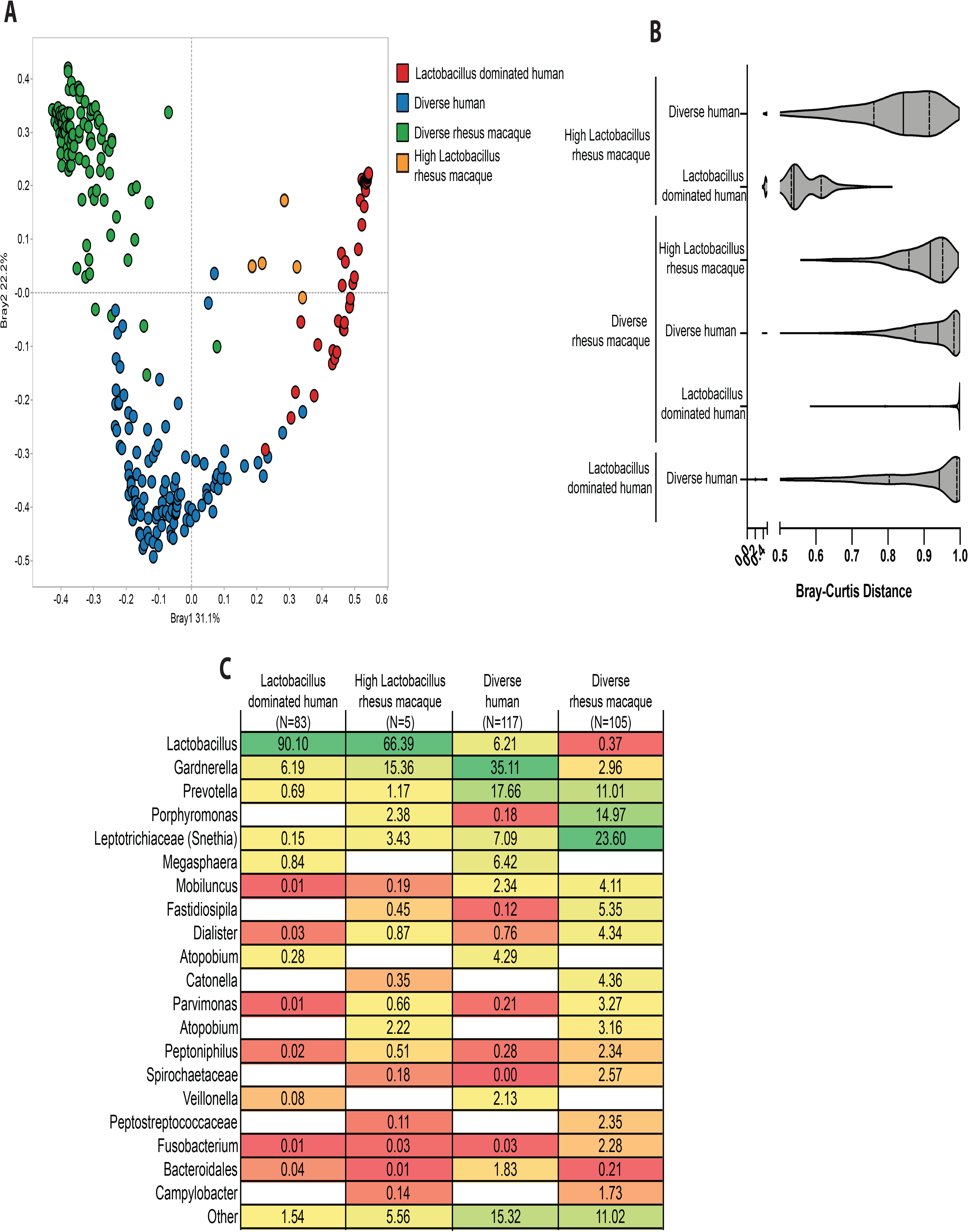
Comparison of human and rhesus macaque vaginal microbiomes. (A) Principal coordinate analysis of Bray-Curtis distance between rhesus macaque vaginal microbiome, Lactobacillus dominated human vaginal microbiome, and diverse non-Lactobacillus dominated human vaginal microbiome samples. (B) Bar graph of the average Bray-Curtis distance between the rhesus macaque vaginal microbiome and the two representative human communities. Significance determined using 1-way ANOVA p < 0.001, with Bonferroni posthoc comparisons, ** p < 0.01, *** p < 0.001. (C) Table of shared and exclusive bacterial genera between the three microbial community types. To be included in this analysis genera had to be present in 10% of samples within a group at < 0.1% relative abundance.

### Metagenomic genome assembly reveals taxa in the rhesus macaque vaginal microbiome similar to human urogenital bacteria

Shotgun metagenomics provides higher resolution taxonomic information and functional potential of microbial communities not attainable using 16S rRNA amplicon sequencing. Shotgun metagenomic libraries were prepared from vaginal samples collected before and 7 days after sucrose/placebo treatment (**Figure 1A**). We eliminated any samples with less than 1 million reads after host decontamination resulting in the loss of 8 samples at the pre-sucrose and 3 samples from the post-sucrose time points (**Supp. Figure 4A**). As noted for 16s rRNA amplicon sequencing, sucrose treatment did not exert a significant impact on the functional potential of the VM (**Supp. Figure 4C**). Although we were unable to identify the *Lactobacillus* colonizing the rhesus VM at the species level using 16S amplicon sequencing or genome assembly, annotation of shotgun metagenomic reads using MetaPhlan revealed the presence of *Lactobacillus johnsonii*, *L. amylovarus*, and *L. acidophilus* (**Supp, Figure 4B**).

We employed metagenomic genome assembly to generate a higher resolution picture of which bacterial taxa were present in the rhesus macaque VM, and how they relate to the genomes of bacterial strains isolated from the human VM. We constructed a total of 78 metagenomically assembled genomes (MAGs) with >80% genome completeness and <2% contamination as measured by CheckM (**Table 2, Supp. Figure 4D**). The MAGs were largely representative of dominant taxa identified in our 16S rRNA amplicon sequencing data, including 9 *Gardnerella*, 6 *Mobiluncus,* 8 *Prevotella*, 2 *Campylobacter* and 3 *Snethia* genomes (**Table 2, Supp. Figure 4D**). We next determined the relationship between our MAGs and human isolates. Macaques *Mobiluncus* and *Snethia* genomes were most closely related to the common human bacteria *Mobiluncus mulieris*, and *Snethia sanguinegens* respectively (**Figure 4 A, B**). Macaque *Gardnerella* genomes were most closely related to *Gardnerella vaginalis*, commonly detected in non-lactobacillus dominated communities and often associated with BV in humans (**Figure 4C**). Additionally, assembled vaginal *Prevotella* and *Campylobacter* genomes were distinct from those we previously assembled from rhesus macaque fecal samples (**Figure 4D, E**). Our assembled *Prevotella* genomes were most closely related to *Prevotella timonensis* and *Prevotella bucallis* previously isolated from human vaginal samples (**Figure 4D**). The assembled *Campylobacter* were most similar to *Campylobacter sputorum* which is most commonly associated with livestock urogenital samples (**Figure 4E**).

**Figure 4:**
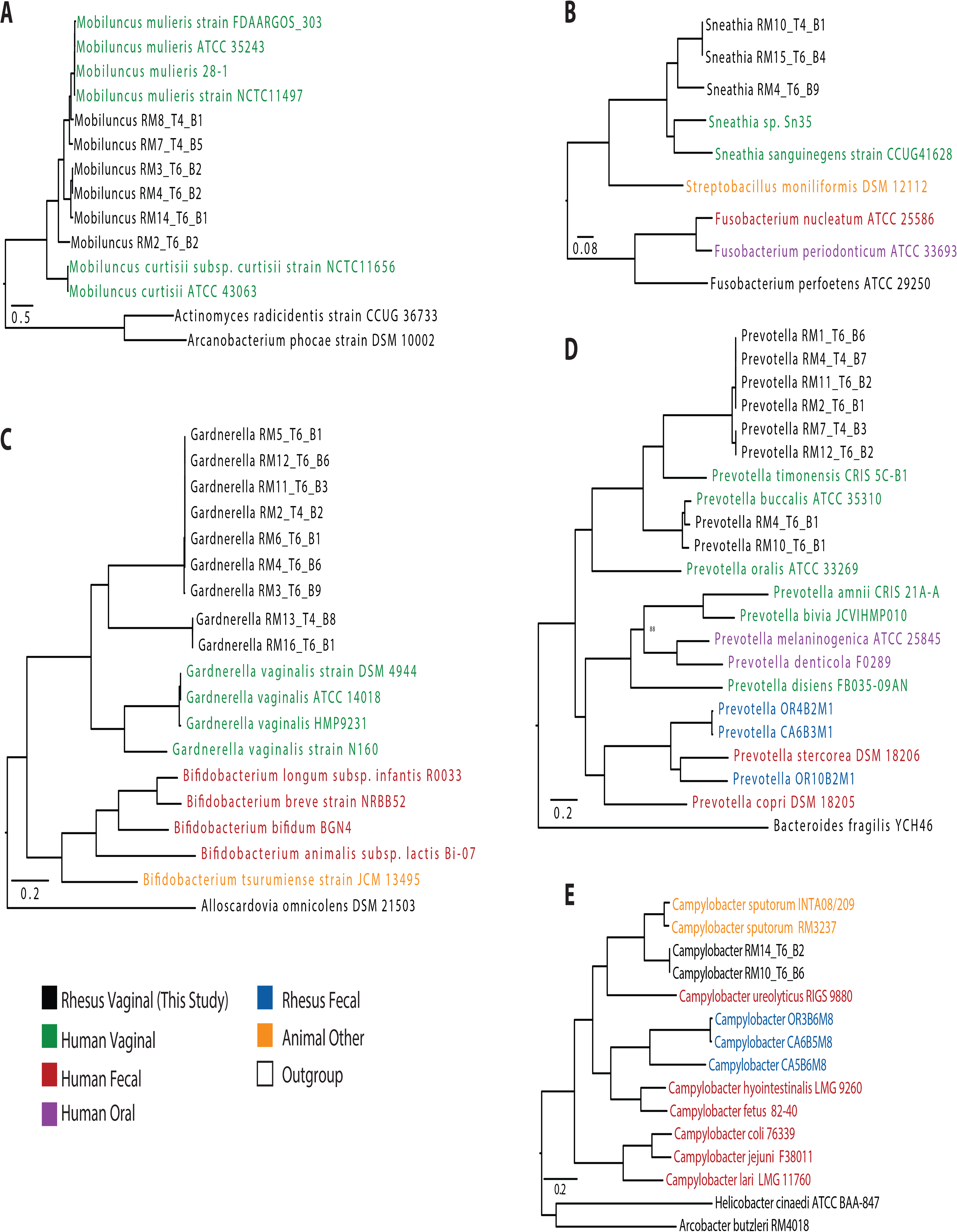
Genomes assembled from the rhesus macaque vaginal microbiome. Phylogenetic tree built using 500 randomly selected conserved Cross-genus gene families (PGfams) for assembled genomes from (A) *Mobiluncus*, (B) *Snethia*, (C) *Gardnerella*, (D) *Prevotella* and (E) *Campylobacter*. Genomes in bold were assembled in this study. Genome ID’s colored by host source.

### The rhesus macaque vaginal microbiome is functionally more similar to women with a diverse vaginal microbiome

To determine the functional potential of the rhesus VM compared to that of human VM, we compared our shotgun metagenomic data to those obtained from clinical studies by Oliver et. al (*46*) and Lev-Sagie et. al (*15*). These studies analyzed samples from human vaginal communities that were classified as “*Lactobacillus* dominated” (>90% *Lactobacillus*), “Diverse asymptomatic” (<90% *Lactobacillus*), and “Recurrent BV”. We used supervised random forest modeling to determine if the overall functional capacity of the VM could be used to distinguish between samples collected from asymptomatic women (either “*Lactobacillus* dominated” or “Diverse Asymptomatic”), women with BV, and rhesus macaques. The overall random forest model was 83% accurate at classifying samples into the four groups with overlap between the VMs from women with BV and asymptomatic women with a diverse VM (**Figure 5A**). The VM of rhesus macaques was more closely related to that of asymptomatic women with diverse communities than to VM of women with BV (**Figure 5B).**

**Figure 5:**
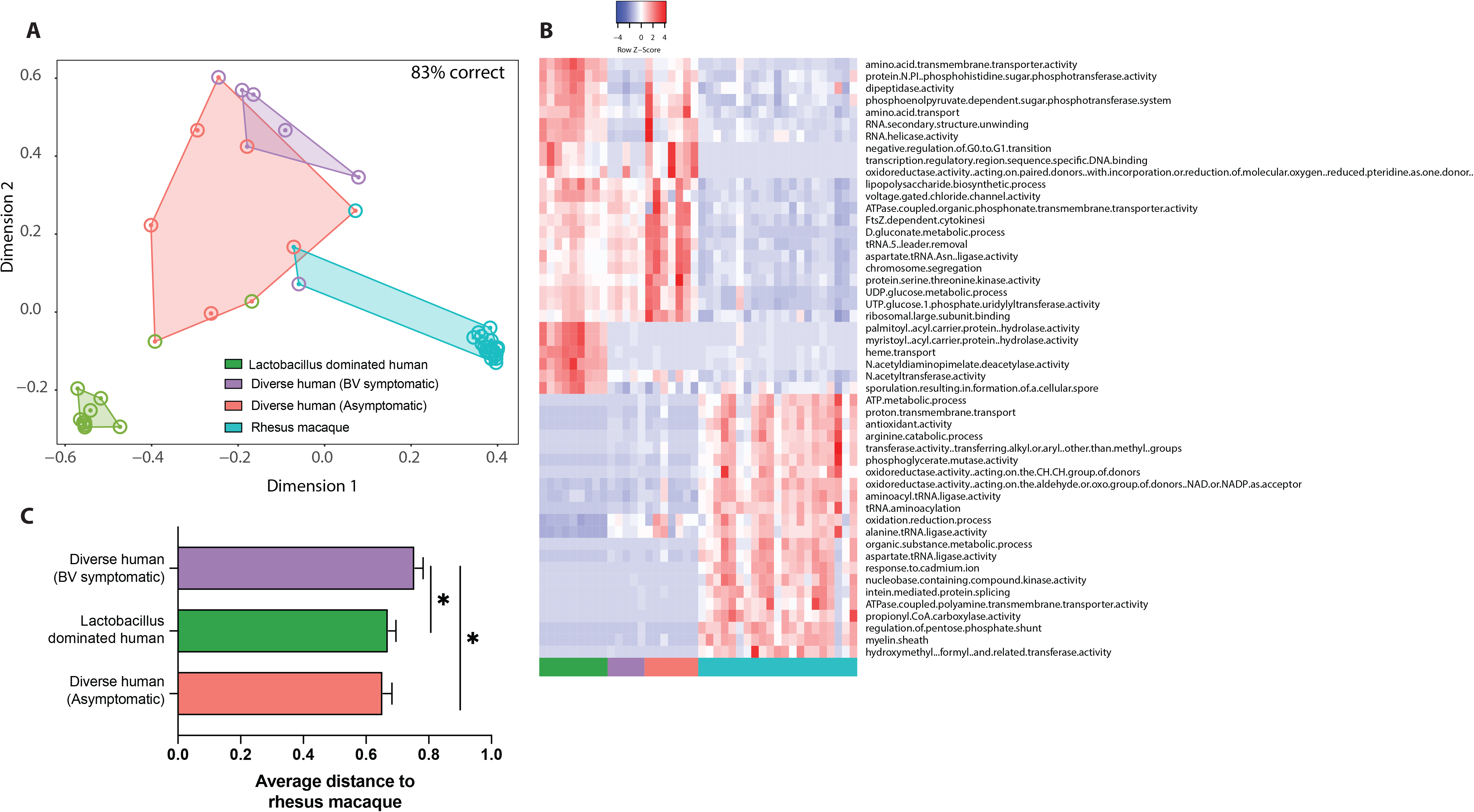
Comparison of the functional potential of the rhesus and human vaginal microbiome. (a) NMDS plot colored by sample source, outer circles denote if a sample was misclassified at any point during the random forest model generation. The color of the outer circle represents which group that sample was misclassified as. (B) Heatmap of 50 most important GO terms as predicted by random forest. (C) Bar graph of the average random forest proximity between the rhesus macaque vaginal microbiome and the three human communities. Significance determined using 1-way ANOVA p < 0.001, with Bonferroni posthoc comparisons, * p < 0.05.

We then extracted the functional GO terms that distinguished between the four groups (**Supp. Figure 4E**). We found that ‘D-gluconate metabolic process’, ‘phosphoglycerate mutase activity’, and ‘lipopolysaccharide biosynthetic process’ were among the most significant GO terms for classifying samples into the four groups (**Supp. Figure 4E**). Several pathways were unique to each group. For example, “palmitoyl-[acyl-carrier-protein] hydrolase activity” and “heme transport” pathways were common in “*Lactobacillus* dominated” human VM communities but absent in vaginal communities from rhesus macaques and from asymptomatic women with diverse communities or those with recurrent BV (**Figure 5C**). Other terms such as ‘D-gluconate metabolic process’ and ‘UDP-glucose metabolic process’ were found in human vaginal communities regardless of health status, but absent in the rhesus VM (**Figure 5C**). We also found GO-terms shared between *Lactobacillus* dominated humans and diverse asymptomatic humans, but absent in women with BV and rhesus macaques such as “protein-N(PI)-phosphohistidine-sugar phosphotransferase activity” and “phosphoenolpyruvate-dependent sugar phosphotransferase system” that were absent in vaginal communities from women with BV and rhesus macaque. We also found that the "oxidation-reduction process” pathway was abundant in rhesus and human diverse vaginal communities but not in *Lactobacillus* dominated human vaginal communities. Finally, we also identified GO terms that were unique to the rhesus macaque vaginal microbiome including “antioxidant activity”, “arginine catabolic process”, and “regulation of pentose phosphate shunt” (**Figure 5C**).

## DISCUSSION

Approximately 50% of women globally have a diverse vaginal microbiome defined by the absence or low abundance of gram-positive *Lactobacillus* that typically dominates the human vaginal microbiome in developed countries (*1, 2, 4, 11*). Instead, this diverse community is composed of gram-negative anaerobic bacteria such as *Gardnerella*, *Prevotella*, *Snethia*, and *Mobiluncus* (*1*). The prevalence of this diverse community can vary widely by geographical location, with a prevalence between 20 and 30% in the developed world (*1, 46*) and between 50-80% in the developing world (*4, 49*). Prevalence of this diverse community state has also been connected to multiple host factors including ethnicity (*50*), genetics (*51*), and sexual habits (*52*). These diverse vaginal communities are largely considered to be a dysbiotic state of the vaginal microbiome, which can result in an inflammatory state such as BV, and increased susceptibility to sexually transmitted diseases (*4, 53*). However, important questions about the implications of this community remain unanswered since many women are asymptomatically colonized by these diverse communities. These questions are challenging to ask in the clinical setting and require the availability of a robust animal model.

In this study, we carried out a comprehensive analysis of the rhesus macaque VM using a combination of 16s rRNA amplicon as well as shotgun metagenomic sequencing. We also measured key clinical parameters associated with BV. Our analysis showed that most animals displayed clinical and microbial markers of BV notably high vaginal pH and high Nugent scores. At the 16S rRNA gene amplicon level, the rhesus VM harbors a diverse set of anaerobic microbes including *Snethia*, *Prevotella*, *Mobiliuncus*, and *Fusobacterium* that are often associated with BV. Although the rhesus macaque VM shares a core set of highly abundant genera with humans, it also harbors a diverse set of anaerobic genera less commonly found in human vaginal samples including; *Porphyromonas*, *Fastidiosipila*, *Catonella*, Peptostreptococcaceae, Spriochaetaceae, and *Campylobacter* (*54, 55*).

Our observations are in line with previous studies that have reported a high prevalence and abundance of these genera in other macaque species (*24–26*). However, our study is the first to report the presence of *Gardnerella*, a key pathogen in human BV, and an indicator of a dysbiotic community (*24–26*). Specifically, *Gardnerella* was present in 62 of 120 samples and at 10 to 66% relative abundance in 15 samples analyzed in this study. Additionally, using shotgun metagenomics, we assembled 9 *Gardnerella* genomes from 9 different animals that were closely related to human *Gardnerella vaginalis*. Moreover, we assembled genomes from a diverse set of vaginal microbes including *Mobiliuncus*, *Sneathia*, and *Prevotella* that were closely related to genomes assembled from the human urogenital tract.

As others have previously reported (*26*), we found that *Lactobacillus* spp. was largely absent in the rhesus macaque vaginal microbiome. In our initial screen, a *Lactobacillus* dominant microbiome was found in 3 of the 10 animals. However, in our longitudinal study, a *Lactobacillus* dominant microbiome (33-87% relative abundance) was only transiently observed in animal RM14 at one time-point and animal RM3 at four time-points. *Lactobacillus* was also detected at between 1-15% relative abundance in 13 samples across 7 animals. In humans, *Lactobacillus* spp. are the major producers of lactic acids that results in low vaginal pH (~3.5). In accordance with those observations, the presence of *Lactobacillus* in these animals correlated with lower vaginal pH and Nugent scores at these time points. Our shotgun metagenomics analysis revealed the presence of *L. amylovarus* and *L. acidophilus*. Although these species have not been identified in samples from women in the developed world, future studies should investigate the presence of these *Lactobacillus* species in VM from women in developing countries.

Previous studies reported that the macaque VM is highly variable over time and that some of this variability was correlated to hormonal cycling (*25*). We observed a similar trend with multiple taxa being correlated to Progesterone levels including *Mobiluncus mulleris* and *Peptoniphilus.* In contrast to previous longitudinal studies, we found that the vaginal microbiome of some animals remained relatively stable while that of others was variable over time. While further studies are needed to determine the causes and consequences of these two unique communities’ states, one potential explanation for this pattern is fecal contamination. Indeed, some samples obtained from 6 of the 9 animals with “variable” communities had fecal contamination while samples from only 1 of 7 animals with “stable” communities exhibited fecal contamination. This may suggest that “variable” VM are responding to community perturbation caused by infiltration of fecal material and microbes. It is also possible that we did not sample often enough to capture the true variability of the rhesus VM. Daily variability has been observed in human samples and was associated with diet, exercise, and natural hormonal cycling (*56*).

We also compared the functional capacity of the rhesus vaginal microbiome of three distinct human vaginal microbiome communities; “Lactobacillus-dominated”, “Asymptomatic diverse” and “Recurrent BV”. This analysis revealed that the rhesus macaque and human VM are functionally distinct. However, the rhesus VM is functionally most similar to that of asymptomatic women with a diverse vaginal community. We found differences in metabolic potential between the four groups with all three human VM functionally enriched in a variety of carbohydrate metabolism GO terms that were not found in the rhesus macaque vaginal microbiome. Host derived glycogens are the major energy source in the vaginal microbiome and driver of *Lactobacillus* dominance (*57*). Differences in host glycogen production between humans and other mammals is hypothesized to contribute to the unique *Lactobacillus* dominated communities found in humans (*7, 58*). Although a small percentage of animals can transiently harbor a *Lactobacillus* dominated VM, microbial communities in rhesus macaques have significantly less vaginal glycogen than humans which may contribute to their inability to sustain a *Lactobacillus* dominated vaginal community (*59*).

We tested an intravaginal prebiotic sucrose intervention to drive the vaginal microbiome to a lactobacillus dominated state and improve clinical markers of BV. Despite previously reported positive results for this intervention (*22, 23*), we found no clinical or microbial indications of improvement. In our hands, intravaginal sucrose gel did not lower vaginal pH, Nugent scores, or Amsel criteria positivity. Additionally, we did not observe an increase in the relative abundance of *Lactobacillus* or a shift in overall community composition. This intervention could have failed because the relative abundance of *Lactobacillus* in these particular animals was below detection at the start of the intervention. Moreover, shotgun metagenomics indicated that the “protein-N(PI)-phosphohistidine-sugar phosphotransferase activity” pathway, a key step in the bacterial import of sucrose, was largely absent in VM of humans with BV and rhesus macaques. A microbial community lacking genes within this pathway is less likely to import and metabolize sucrose. It is also worth noting that since the original goal of this study was to determine if intravaginal sucrose treatment could improve clinical markers and increase the abundance of *Lactobacillus,* we excluded animals with low vaginal pH and high *Lactobacillus* abundance. In future studies, these animals should be identified and studied longitudinally to determine if this represents a stable or transient community state.

It has been well established that humans in the developing world have a distinct microbiome from those of the developed nations (*60*). This has been particularly well characterized in the context of the gut microbiome. For example, individuals in the developing world harbor a more diverse gut microbiome colonized by microbes that have all but disappeared in the western microbiome (*61*). This is due to a combination of key environmental factors such diet, and antibiotic use among other factors (*62–64*). In humans, there is also a clear distinction between the vaginal microbiomes of individuals of the developed and developing worlds. Specifically, women in the developing world have a much higher prevalence of diverse non-*Lactobacillus* dominated communities (*4, 49*). Understanding the drivers of these distinct microbial community states is critical to developing VM targeted therapies. As we have previously shown for the gut microbiome, we found that the rhesus VM is also more reflective of women in the developing world (*30*).

In summary, we show that macaques harbor a diverse vaginal microbial community similar to that detected in women with a non-*Lactobacillus* dominated community or symptomatic BV at the 16S rRNA gene amplicon level. Additionally, we show genomes assembled from the rhesus macaque vaginal microbiome are closely related to human pathobionts associated with BV. Future studies should focus on the immunological landscape of the rhesus urogenital tract to determine if this diverse community state results in local inflammation as seen in humans with BV and asymptomatic women with diverse communities. There are many unanswered questions about why this community is maintained in some women and populations. While this community type has been shown to increase the risk for some sexually transmitted diseases such as HIV and HPV, these communities may be important for defense against pathogens that are more prevalent in the developing world. Rhesus macaques offer a unique opportunity to explore the importance of a diverse vaginal community that is highly prevalent in women on a global scale.

---

**Supplemental Figure 1: Removal of vaginal samples with potential fecal contamination.** (A) Principal coordinate analysis of unweighted UniFrac distance generated from vaginal and concurrently collected fecal samples colored by sample source. Vaginal samples circled in black denote samples with suspected fecal contamination. (B) Scatter plot of measured absolute sequence variants (ASVs) for each sample type including vaginal samples with suspected sequence contamination. Significance determined using 1-way ANOVA p < 0.001, with Bonferroni posthoc comparisons, *** p < 0.001. (C) Stacked bar plot highlighting three taxa that were shared between fecal samples and vaginal samples with fecal contamination but absent in all other vaginal samples and the sequenced microbial community standard. (D) Table of all vaginal samples used for 16S rRNA gene amplicon sequencing. Boxes highlighted in red were samples eliminated due to suspected fecal contamination, boxes in white were eliminated from alpha and beta-diversity analysis due to low sequencing depth. Boxes in green denote samples that were retained for further analysis.

**Supplemental Figure 2: Impact of intravaginal sucrose treatment on clinical markers of BV in rhesus macaques.** (A-F) Longitudinal measurements of (A) Nugent scores, (B) vaginal pH, (C) Whiff positivity, (D) clue cell positivity, (E) levels of estradiol, and (G) progesterone.

**Supplemental Figure 3: Impact of intravaginal sucrose treatment on the vaginal microbiome in rhesus macaques.** (A) Principal coordinate analysis of weighted UniFrac distance colored by treatment group at baseline (open circle) and 14-days post-treatment (filled circle). (B, C) Longitudinal measurement of (B) ASVs and (C) Shannon evenness divided by treatment group. (D) The relative abundance of Lactobacillus separated by treatment group at baseline and 4 time-points post-treatment.

**Supplemental Figure 4: Shotgun metagenomic analysis of rhesus macaque vaginal microbiome.** (A) Scatter plot of sequences remaining after removal of host reads. Red indicates samples that were removed from downstream analysis due to low sequencing depth. (B) Table of genomes assembled based on assigned genus. (C) NMDS of shotgun metagenomic samples collected 7-days post sucrose treatment colored by treatment group, outer circles denote if a sample was misclassified at any point during the random forest model generation. The color of the outer circle indicates which group that sample was misclassified. (D) Heat map of top 50 GO terms ordered by the overall contribution to random forest prediction accuracy, colored by importance for defining each group and their contribution to overall model accuracy.

